# Long time-scales in primate amygdala neurons support aversive learning

**DOI:** 10.1101/263889

**Authors:** Aryeh H. Taub, Tamar Stolero, Uri Livneh, Yossi Shohat, Rony Paz

## Abstract

Associative learning forms when there is temporal relationship between a stimulus and a reinforcer, yet the inter-trial-interval (ITI), which is usually much longer than the stimulus-reinforcer-interval, contributes to learning-rate and memory strength. The neural mechanisms that enable maintenance of time between trials remain unknown, and it is unclear if the amygdala can support time scales at the order of dozens of seconds. We show that the ITI indeed modulates rate and strength of aversive-learning, and that single-units in the primate amygdala and dorsal-anterior-cingulate-cortex signal confined periods within the ITI, strengthen this coding during acquisition of aversive-associations, and diminish during extinction. Additionally, pairs of amygdala-cingulate neurons synchronize during specific periods suggesting a shared circuit that maintains the long temporal gap. The results extend the known roles of this circuit and suggest a mechanism that maintains trial-structure and temporal-contingencies for learning. It further suggests a novel model for maladaptive behaviors.

## Introduction

In associative learning, the passage of time between trials – the inter-trial-interval (ITI), can potentially serve as a cue of trial expectation. The time that passes from the offset of one trial carries information about the onset of the next trial, and If this time can be internally represented and kept, it can aid to form higher-order representations of the environment. Evidence that the ITI is indeed used for timing and auto-shaping of responses was first described in a series of classical work^1–3^. Importantly, the ITI length affects acquisition rate and memory strength^4–6^, and ITI-to-ISI ratio predicts acquisition characteristics^7, 8^. This suggests that acquisition of associative memory involves assessment and integration of multiple temporal contingencies, and specifically, that temporal information about the ITI is internally represented even in the absence of CR-like behavior during it.

Because the ITI usually lasts from few seconds to dozens of seconds, it raises the question where is this temporal lag being maintained? Neurons in the basolateral complex of the amygdala (BLA) play a role in acquisition and expression of affective associations ^9–12^, and show tonic responses and baseline changes that last after a specific trial or stimulus end^13–15^. The amygdala also plays a role in the acquisition of trace-conditioning, where a temporal gap between the conditioned stimulus (CS) and the unconditioned stimulus (US) must be bridged^16–20^, and even in longer lags that were traditionally thought to be dependent on the hippocampus^21, 22^. In addition, the amygdala plays a role in 2^nd^-order conditioning^23–25^, that can be formed between the ITI itself and the CS^26^; and even hold abstract representations of the trial structure beyond the CS-US relationship^27, 28^. Further, the Amygdala can use its bi-directional connectivity with the anterior-cingulate-cortex (ACC)^29^. The ACC is required for trace-conditioning with long gaps^30^ likely due to its role in attention^30–32^, it forms and integrates representations of task structure^33, 34^, and exchange information with the amygdala for flexible updating of contingencies^19, 20, 35, 36^.

We therefore hypothesized that the BLA together with the ACC hold a representation of long-time scales during the ITI of an aversive conditioning task, even without an explicit external cue. We further hypothesized that ITI representation contributes to the formation of the conditioned responses (CRs) without behavioral manifestation during the ITI itself. To address this, we recorded the activity of single neurons in the amygdala and the dorsal-ACC (dACC) of non-human primates during a tone-odor conditioning paradigm with a relatively long (dozens of seconds) ITI that varies on a trial-by-trial basis.

## Results

We recorded neuronal activity during the inter-trial-interval (ITI) in a tone (CS)-odor (US) conditioning task, where an aversive odor (propionic-acid) was paired with a pure-frequency tone (250 ms, randomly chosen each session from 900-2400 Hz). Tones were triggered by onsets of respiration cycle detected in real-time, and odors were presented at the onset of the following respiration cycle (mean ISI duration of 1.6 ± 0.24 sec). As shown in our previous studies^19^ and here (Fig. 1A, B), pairing resulted in a CS-evoked augmented inhale that preceded the presentation of the odor, and reflects a learned preparatory response (CR).

**Figure 1.**
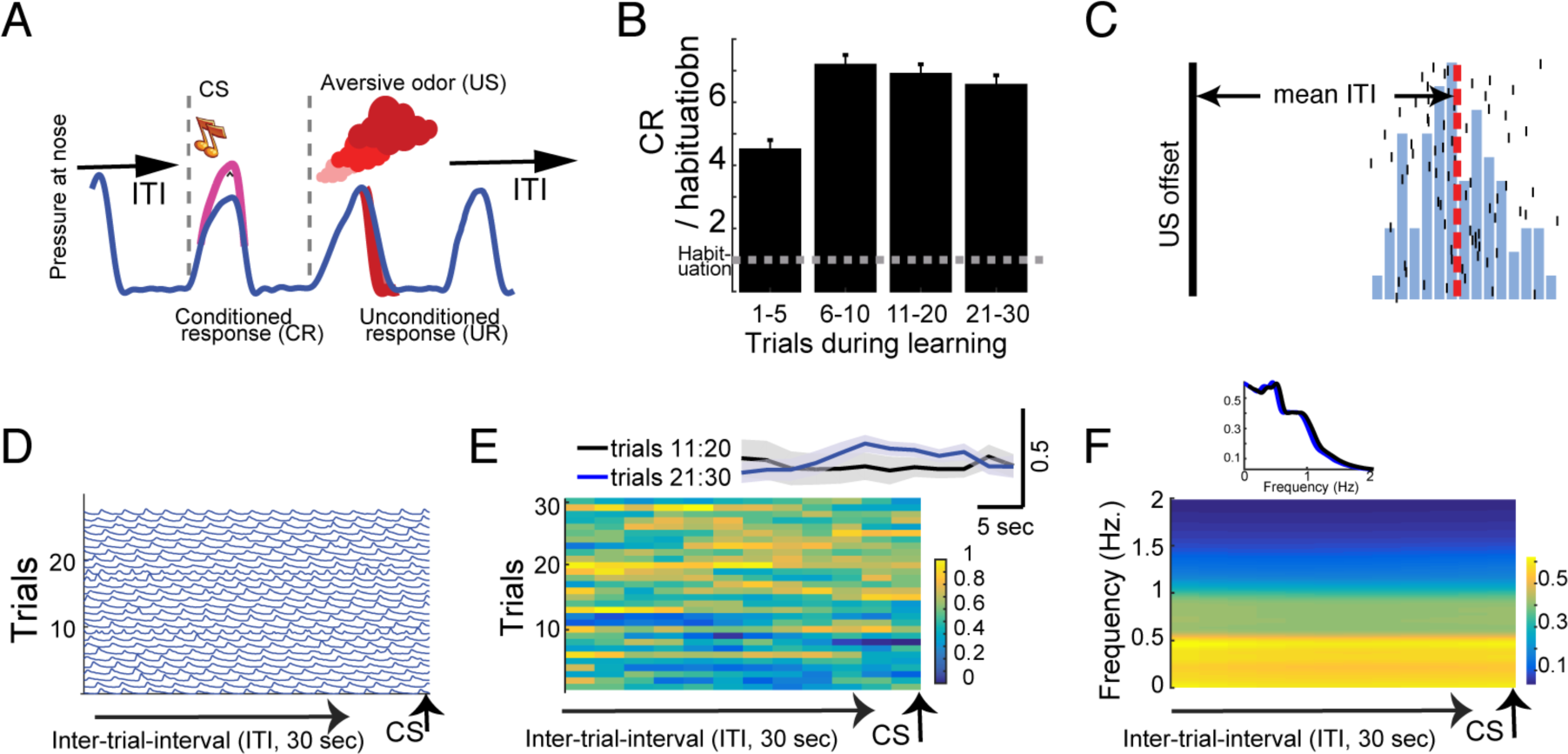
Conditioning paradigm and behavior during the inter-trial-interval (ITI) (A) Online detection of inhale is used to initiate tone(CS) presentation that is followed by an aversive odor(US) at the onset of the next inhale. Response to the aversive odor is a decreased inhale (UR), whereas the conditioned response (CR) is an increased inhale. (B) The CR increases during learning. Shown is the CR (inhale-size during the 350 ms post-CS onset) as proportion of the inhale-size during habituation (dashed line). (C) A histogram of ITI duration from one representative session. Trial occurrences (black vertical line signals US offset) and the initiation of the subsequent trial (black vertical ticks). Dashed red line is the mean ITI at 42 seconds. (D) Inhale traces during the ITI aligned to onset of next trial. Shown is data for one session of learning. No clear modulation can be observed towards the end of the ITI. (E) Inhale modulation during the ITI did not change across learning. Shown is mean across all learning sessions for the 30 learning trials (one-way ANOVA, p > 0.1, df=11, f=0.05). Inset presents mean over trials 11-20 and 21-30, computed during 30 sec. of ITI. (F) No change in inhale frequency across learning trials. Shown is mean power across all learning sessions at each frequency (0.2 – 2 Hz.) during the ITI (one-way ANOVA, p<0.01, df=11, f=0.05). Inset presents inhale frequency (mean over days) as power during the whole 30 sec. of ITI preceding the CS, separately for trials 11-20 and 21-30.

### ITI length contributes to learning-rate

As can be expected from the long ITI duration and its unreliable trial-by-trial timing (mean ITI 38.7 ± 4.4 sec, Fig.1C), combined with the fact that the following CS was present before the US (i.e. a complete predictor), we did not observe any preparatory behavioral response during the ITI (Fig.1D), neither in the depth of inhale modulation (Fig. 1E) nor in its frequency (Fig. 1F).

However, when sorting the CR based on the duration of the previous trial ITI, we found that the length of the ITI induced a higher CR, both in absolute value compared to habituation (Fig. 2A, B), and in increased learning rate (Fig. 2C). These results show that longer ITIs induced more learning-specific behavioral response.

**Figure 2.**
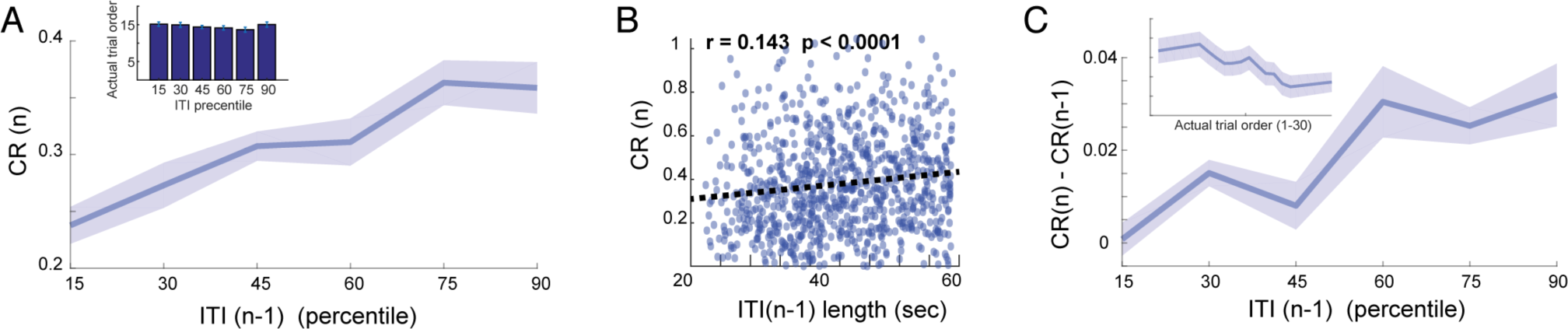
Length of ITI contributes to learning-rate. (A) The size of the CR in a trial (n) as a function of ITI duration in the previous trial (n-1), averaged across all sessions and binned into 6 percentiles (one-way ANOVA, significant effect for percentile, df = 5, f = 29.85, p < 0.0001). Inset shows the averaged trial order sorted similarly by ITI duration percentile (n = 171, one-way ANOVA, df = 5, f = 0.74, p = 0.59). (B) Furthermore, CR size was positively correlated with the previous ITI duration (Spearman correlation, r = 0.14, p < 0.0001) on a trial-by-trial basis. (C) Change in CR amplitude between successive trials (CR in trial(n) minus CR in trial(n-1) increase with ITI duration (one-way ANOVA, df = 5, f = 81.14, p < 0.0001). ITI duration (x-axis) computed similarly to (A). For control, Inset shows the standard expected learning effect i.e. the same change in CR(n) – CR(n-1) as a function of trials in learning.

### Neurons modulate their activity during the ITI

We recorded single-unit activity in the amygdala and the dACC (Fig. 3A, amygdala: n = 291; dACC: n = 263), and first validated previous results^19^ by comparing pre-CS to post-CS activity (paired t-tests, p < 0.05). We found that 24% (n=70) of amygdala and 29% (n=76) of dACC neurons had CS-evoked activity, confirming learning related changes in these regions (p<0.001, binomial test).

**Figure 3.**
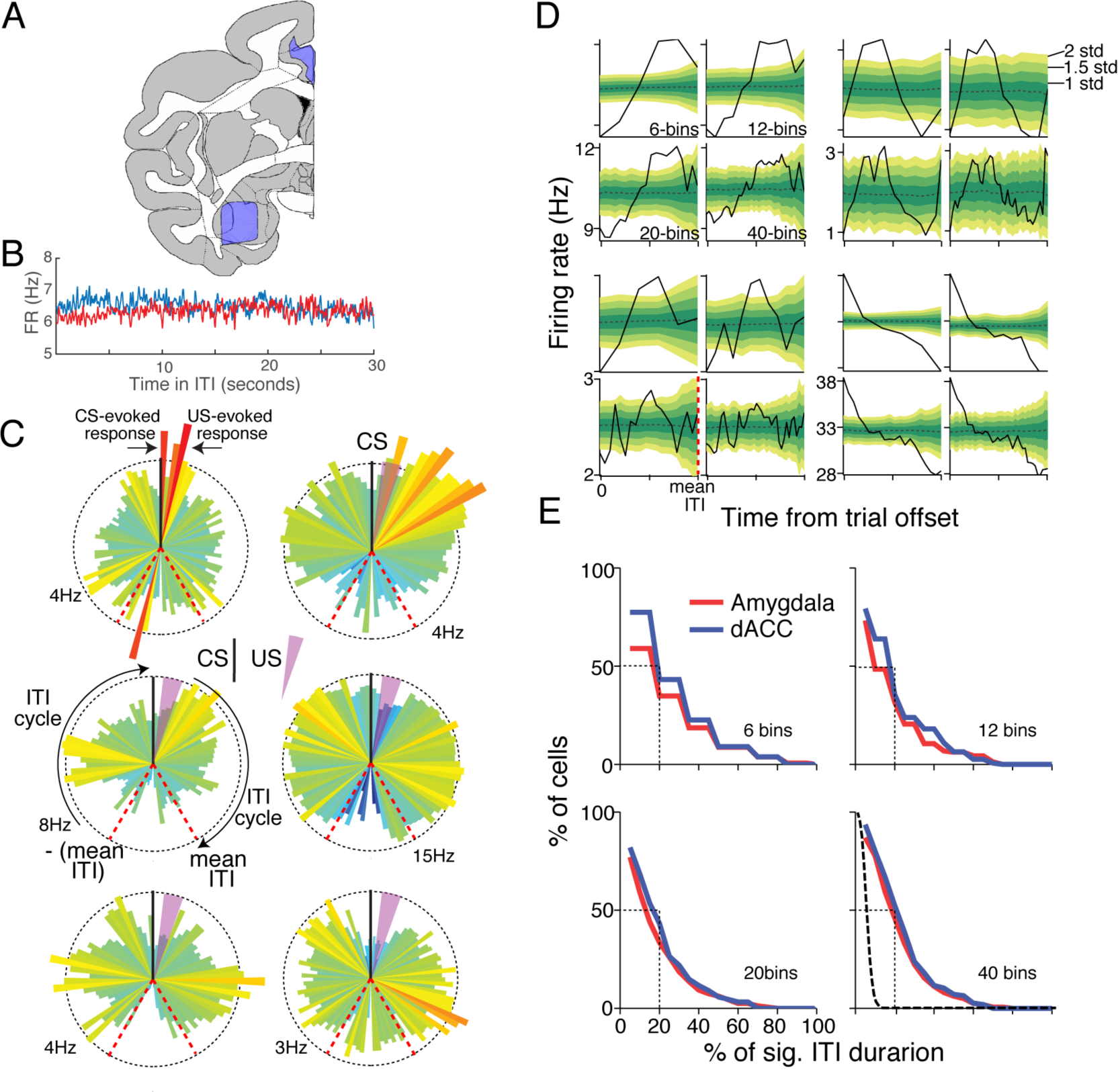
Neurons in the amygdala and the dACC are modulated during the ITI. (A) Recording locations were all made within the purple surfaces, shown on top of a macaque atlas (at AC=0, IA=+20mm). (B) Firing rate (FR) during 30 seconds aligned to next trial CS, averaged over all sessions and neurons in both Amygdala and dACC. No period showed specific modulation. (C) Single-cell firing-rates of 3 Amygdala neurons (left column) and 3 dACC neurons (right column). Modulation occurs not only in response to the CS and the US (as in the left top example), but also during different periods in the ITI. Shown are PSTHs for 2 ITI cycles, before and after CS-US (CS is marked by black vertical line at 90 ° and US by light purple wedge). Cells exhibit temporal-tuning in different periods during the ITI. (D) ITI-FR modulation evaluated from trial offset (time 0) to the mean ITI (dashed red line). Shown are 4 single-cell examples of the actual ITI-FR (black line) binned into 6, 12, 20 or 40 bins (for robustness) and the shuffled confidence interval (yellow-green shades). Notice that data was shuffled only between bins that occur before a new trial initiates, and as a result the confidence interval increases towards the end of the ITI (E) Histograms of the proportion of cells (y-axis) that showed significant ITI-FR modulation (ITI-FR > 2 s.t.d of the shuffled FR) in a proportion of the ITI duration (x-axis). For example, 50% of the cells had significant modulation in 20% or more of the ITI duration were (dashed black line). All binning options are shown for robustness. A null distribution (obtained from shuffling) is shown for comparison in dashed black line, where most cells had significant modulation at around 5% of the time (as expected from chance).

We next turned to examine the activity during the ITI. On average, no specific epoch of the ITI showed modulation of activity (Fig. 3B). However, close inspection of the data suggested that many of the neurons do modulate their firing rates during the ITI, but over specific temporal scales (Fig. 3C). To test this, we analyzed the neuronal activity between the offset of each trial and the expected initiation of the next trial, namely the average ITI length (‘mean-ITI’), computed separately for each session. We omitted the bins where the CS occurred before the mean-ITI to avoid CS-evoked activity in the analyses. Neurons’ discharge during the ITI were binned into segments, and the strength of the ITI modulation was assessed by comparing the original ITI-firing rate (FR) with shuffled data (Fig. 3D). Next, we determined for each bin independently if its firing rate differs from the shuffled data. We found that the activity of roughly 50% of amygdala and dACC cells differed from baseline activity for 20% or more of the total duration of the ITI (Fig. 3E, p < 0.01, binomial test). For robustness, we tested and observed identical results under different segmentations (6, 12, 20 and 40 bins, Fig. 3D, E). For comparison, a null distribution derived from shuffled data showed that almost all cells have significant activity in < 5% of ITI duration, as expected from chance-level modulation (Fig.3E). We conclude that in both structures, large and similar proportions (p > 0.1, binomial tests) of cells had significant modulation during the ITI (see also Supp. Fig.1).

### ITI modulation strengthens during conditioning

If indeed ITI modulation in single cells is related to the acquisition process, then we can expect it to strengthen during learning. To evaluate the strength of the ITI-FR modulation we computed the excess in spikes after normalization to the mean and standard deviation of the shuffled data, to allow comparison across cells (Fig. 4A, B). The root-mean-square (RMS) of 21% (n=61) of amygdala neurons and 26% (n=68) of dACC neurons was increased during one or more stages (early, mid, or late, 10 trials each) of the acquisition compared to habituation (one-way ANOVA, p < 0.05). These acquisition responsive neurons (23%, 129/554) were further inspected during same-day extinction that started 15-30 minutes after the acquisition ended. We found that their ITI modulation diminished significantly yet remained higher than habituation (one-way ANOVA, p < 0.01 for both regions, Fig. 4B). This result suggests that during acquisition of emotional memory, the activity of neurons in the amygdala and the dACC becomes locked to cycles of CS-US and the long ITI (dozens of seconds) between them.

**Figure 4.**
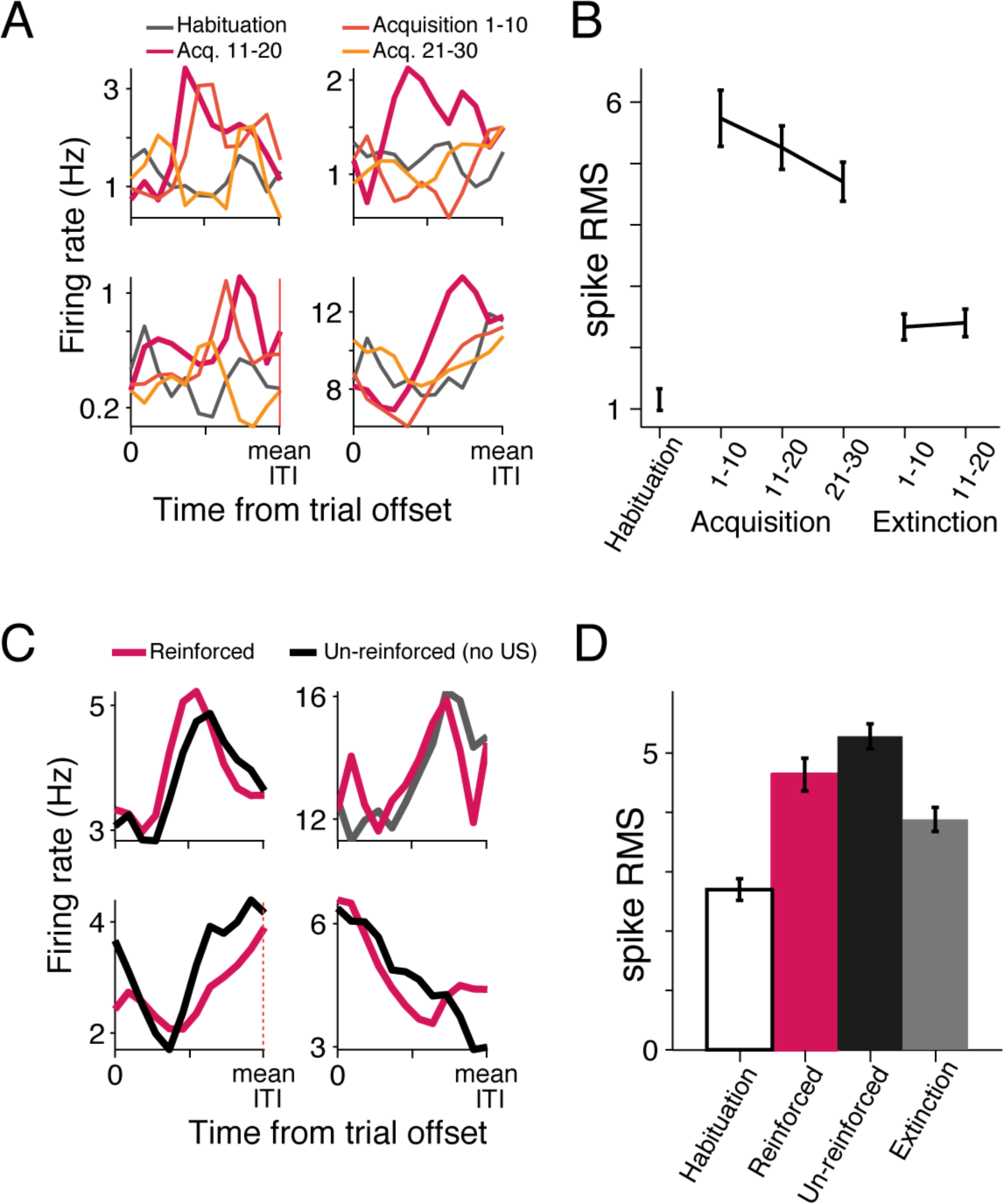
Modulation in ITI-FR increases during acquisition of emotional memory. (A) Shown are four examples of single-cells with ITI-FR modulated more strongly during the acquisition (yellow-red shades) than during habituation. (B) The mean difference in spikes during the ITI-FR (over shuffled data, quantified as RMS), increased during acquisition, and diminished partially during extinction. Shown are the mean values for all cells with significant modulation above habituation (see text and methods). (C) ITI-FR modulations in un-reinforced trials (no US) was similar to the ITI-FR modulation in reinforced trials. Shown are 4 single-cell examples. (D) Cells with ITI-FR modulation in un-reinforced trials had similar modulation for the reinforced trials, and reduced ITI modulation in extinction (average over all cells with significant modulation during un-reinforced trials).

Although odor exposure can remain in the receptors for few seconds and even induce prolonged behavioral effect, it cannot account for the ITI modulation we observe. First, the neural modulation often initiated during the middle and late segments of the ITI (Fig. 3, and see next sections), yet to completely exclude this possibility we introduced *‘catch’* trials with un-reinforced CS during the acquisition (< 1/3 of the trials, during half of the sessions). We found that 11% (29/247; p<0.05, binomial test) of the cells had significant modulation in the ITI following un-reinforced trials (notice that the lower number of cells can stem from the lower number of trials used for analysis, and hence lower statistical power). Importantly, the ITI response of these cells was highly similar in reinforced and un-reinforced trials (Fig. 4C, D, t-test, p > 0.1). Here again, their response diminished during extinction (Fig. 4D; t-test, p < 0.05). In sessions that included both aversive and appetitive odors conditioned to two different tones (discrimination learning), we found that the proportion of ITI-FR modulation was significant after both types of valence, but higher following aversive trials (Supp. Fig. 1).

Overall, the findings are in line with the hypothesis that the FR modulations during the ITI contribute to learning by signaling ITI length.

### Trial-by-trial modulation of ITI signaling

If indeed expectation is formed based on the mean of the inter-trial-interval, and ITI length contributes to learning-rate (Fig. 2), one can hypothesize that slight variations in length can induce transient updates in representation. Specifically, the duration of the ITI at the previous trial (n-1) is expected to correlate with the time of modulation (peak-activity) during the following trial (n). To explore this possibility, correlations were calculated between the time of modulation (center-of-mass for the ITI-FR) in the current trial and the duration of the ITI of the previous trial, for all available neurons. Indeed, 15% of the neurons significantly correlated with the duration of the previous ITI (Fig. 5A, B). Interestingly, correlations were significantly more common in dACC neurons than in amygdala neurons (17.4% and 10.6%, respectively, p<0.01, χ2 test for indepencdence), yet in both regions their prevalence exceeded chance-level (p < 0.05, binomial test). Earlier ITIs (of trials n-2, n-3 and n-4) had only limited effect on the center-of-mass (7-9% of cells, p > 0.05), and a multiple regression model including n-1/2/3/4 suggested that the combined duration affect only few cells (4%, data not shown). This result demonstrates that the neural activity during the ITI is constantly updating by the most recent trial.

**Figure 5.**
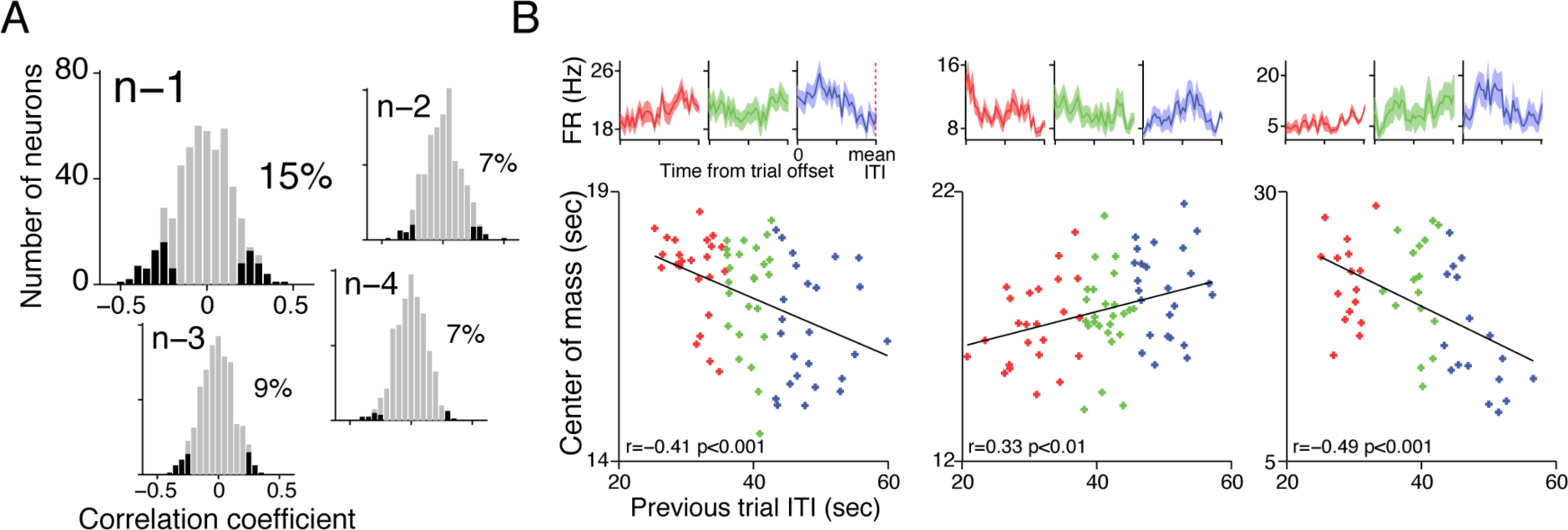
ITI duration affects the preferred-time of temporal-tuning in the next trial. (A) The location of the preferred-time is correlated with the duration of the previous trial ITI in 15% of all cells (p<0.01, binomial test). Earlier ITIs (trials n-2/3/4) affect much less. (B) Three examples of cells in which the preferred-time (quantified as center-of-mass, y-axis) correlates with the ITI duration of the previous trial (x-axis). The PSTHs of the lower third (red), middle third (green) and upper third (blue) are shown on top.

In addition to modulations in peak-timing, we identified a subgroup of neurons in which the strength of the modulation was correlated with length of the ITI in the previous trial; that is, a relatively long ITI in the previous trial induced a stronger modulation during the ITI of the next trial (Supp. Fig.2, 15% of neurons, equally distributed in the amygdala and the dACC). This further demonstrates that neurons adapt their neuronal tuning during the ITI based on the duration of the previously encountered period.

### Amygdala and dACC possess temporal-tuning but signal different periods

Our results so far indicate that amygdala and dACC neurons modulate their activity during the ITI, but is it a neural representation that spans the complete ITI duration? To address this, we inspected the distribution of peak locations of the modulation of amygdala and dACC neurons. Overall, 35% of amygdala cells (101/291) and 33% of dACC cells (89/263) exhibited a temporal tuning-curve with distinct peak (“preferred-time”, methods).

Surprisingly, although the proportion of amygdala and dACC cells with significant tuning modulation was highly similar (p > 0.1, binomial test), the distributions of peak locations were substantially different (Fig. 6A, B). Whereas amygdala neurons had a higher density of peaks at the center of the ITI, the distribution of dACC peaks was bimodal, with most cells modulating their rate either early or late in the ITI (Fig. 6A, B, p < 0.05, permutation test comparing the regional distributions). The mean width of the peaks was similar in the amygdala and the dACC (p > 0.01, t-test), and in both regions the width of the peaks were narrower at ITI margins (Fig. 6C; p < 0.01, one-way ANOVA). These results suggest that the amygdala and the dACC signal different periods within the ITI, potentially enabling maintenance of time over large-scales in a shared network.

**Figure 6.**
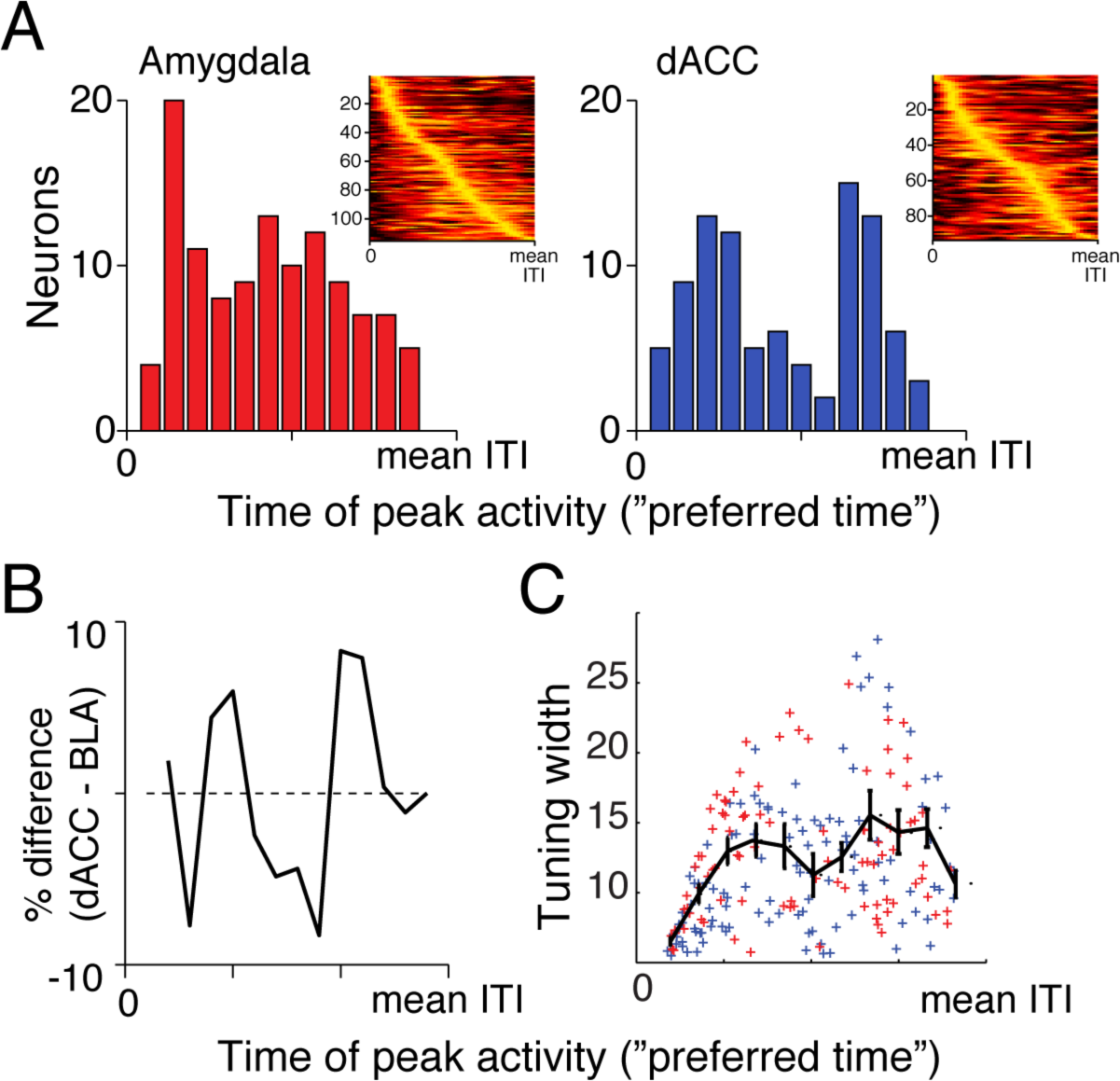
The peaks of temporal-tuning in the amygdala and dACC are differentially distributed. (A) Histograms of the locations of ITI-FR peaks within the full duration of the ITI. Amygdala (left) peaks are uniformly distributed with a slight preference to peak at the middle part of the ITI (the initial narrow peak is likely related to US response). dACC (right), on the other hand, mostly peaked at the early and late phases of the ITI. Insets show normalized PSTHs of all cells with significant modulation during the ITI, sorted by peak location. (B) The difference between these two histograms (p<0.05; permutation test between distributions). (C) The tuning-width (half-width at half-height around the peak) was slightly narrower at edges of the ITI, but maintained an otherwise homogenous width in both regions. Single points mark individual neurons in amygdala and dACC, black line indicates the mean and SEM over all neurons.

### Synchronized amygdala-dACC activity communicates temporal information

If amygdala and dACC represent different types of information and time during the ITI, it is possible that they transfer and share such information to aid in the maintenance of long time scales. To examine this, we computed inter-regional correlations between pairs of amygdala and dACC neurons that were recorded simultaneously. The cross-correlations were computed in 3-seconds window that advanced in 1-second steps from the end of each trial.

Evaluation of significance in such cross-correlations requires two sets of shuffled data to address two different null-hypotheses (Supp. Fig. 3). First, data was shuffled *within-ITI,* so that spikes from a specific ITI were shuffled within itself, thus maintaining the overall firing rate in each ITI. This shuffle targets pseudo-correlations that may arise when the overall ITI-FR rate of the two units covaries. Second, data was shuffled *across-ITI*, so that spikes are shuffled between different trials’ ITI - but from the same period within the ITIs. This addresses pseudo-correlations that may originate when the ITI-FR is repeatedly modulated along the ITI (Supp. Fig. 3). Only cross-correlations that differed from both were deemed significant. A total of 41% (344/839) pairs of amygdala-dACC neurons were found to have a significant cross-correlation. Most correlations had a zero-lag with no clear physiological direction, suggesting reciprocal interaction between the amygdala and the dACC (Fig. 7A). However, the distribution of significant cross-correlations peaked mainly at early and late periods of the ITI (Fig. 7A).

**Figure 7.**
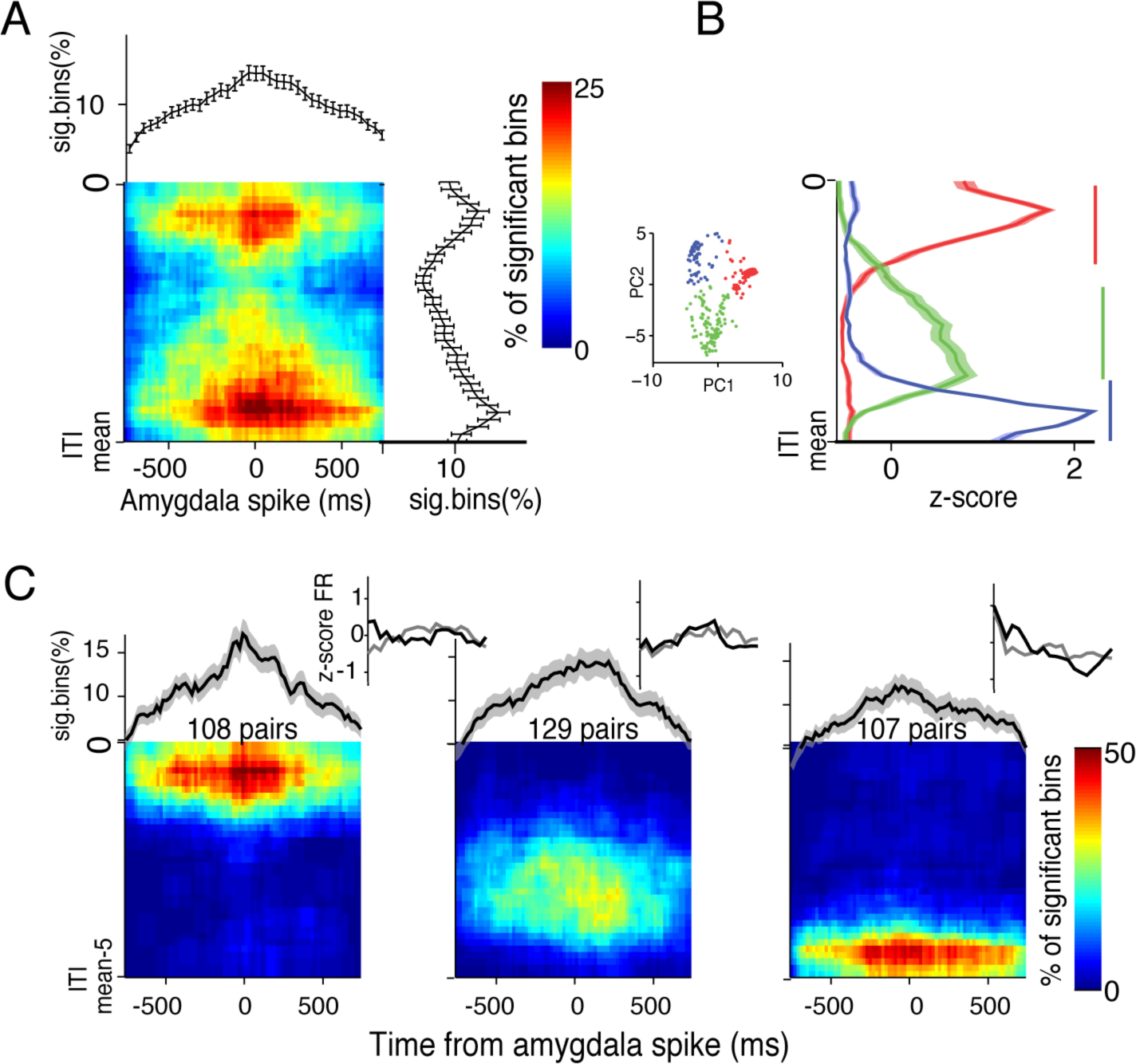
Amygdala-dACC pairwise-correlations peak at specific times during the ITI. (A) Cross-correlations were computed between simultaneously recorded amygdala-dACC neurons in sliding 3-seconds segments of the ITI. CCs were compared to shuffles to exclude changes resulting from covariations within and across ITIs (methods and Supp.Fig.3). Shown is the map of % significant bins from all pairs, for the duration of the ITI (y-axis, top-to-bottom) and triggered on Amygdala spike (x-axis, time zero). Upper plot shows the average (marginal) CC over the whole ITI, and right plot shows the average (marginal) percentage of sig. bins along the ITI. (B) Shapes of individual CCs were clustered (inset, k-means on principal components after z-score), and the average shape for each cluster is presented; revealing two distinct times of CC peaks – early (red) and late (blue) in the ITI, and a 3^rd^ (green) lower widespread modulation. Horizontal colored bars indicate regions with significant synchronization (p<0.01; t-tests). (C) Same as in (A) presented separately for each cluster found in (B). Pairs with early (left) and late (right) CC were more focused in time during the ITI and exhibited stronger synchronization.

We next examined if this double-peaked pattern reflects an average correlation map of prototypic pairs of neurons, or a mixture of single-peaked correlations. A principle component analysis (PCA) on the normalized distribution of correlation density along the ITI identified 3 separable clusters (Fig. 7B, C; k-means clustering). The distribution of cross-correlations of each cluster strengthens the previous finding and shows that it is comprised of different pairs that synchronize during a single and confined segment of time within the ITI. These segments tile the complete duration on one hand (Fig. 7C), but are much more dominant early and late during the ITI.

The results suggest that BLA-dACC neurons signal and transfer temporal information in a pairwise-specific manner. It is also in-line with the aforementioned finding of differential modes of temporal-tuning, and suggests that transfer of temporal information occurs between the regions to enable the differential modes of peak distributions in the two regions.

### Estimating time during the ITI with neural population

If neurons have temporal-tuning with peaks in different times during the ITI, and these are relatively homogenously distributed when combined across the amygdala and the dACC, then one should be able to decode the specific time during the ITI (and as a result, its total length as well). To test this, we used an optimal-linear-estimator (OLE) with cross validation, and found that time could be estimated based on the population activity within a reasonable error (Fig. 8A), that dropped with the size of the population being used, yet remained within few (<4) seconds accuracy (Fig. 8B). Notice this is an underestimate because we assume independence between neurons (similar results were obtained with a population-vector-approach and a Naïve Bayes classifier). Averaging the estimates showed that it is unbiased (Fig. 8C), and therefore a downstream region can use this information reliably to assess ITI duration.

**Figure 8.**
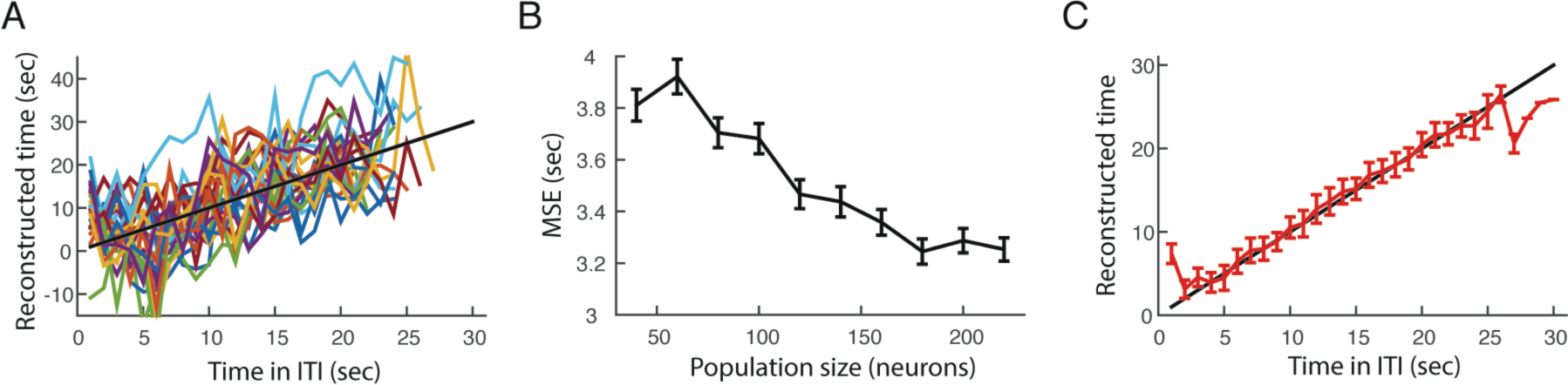
Time in ITI obtained from neural activity. (A) Decoding the time in the ITI from neural activity. Shown are 25 individual repetitions, when calculating each neuron preferred-time on a set of 25 trials (drawn pseudorandomly for each repetition), and using it to estimate time from the actual firing rate in the remaining 5 trials. Black line marks identity line (perfect prediction). (B) The estimation-error as a function of number of neurons used (neurons were taken as independent and collected from all days). The error drops, but seems to plateau at 3 seconds resolution. (C) Average of 100 repetitions using all available neurons. The population estimates the time in the ITI with very small bias (edges are biased due to floor/ceiling effects). Black line marks identity line (perfect prediction).

## Discussion

We show here that neurons in the basolateral complex (BLA) of the primate amygdala together with neurons in the dorsal-anterior-cingulate-cortex (dACC) acquire temporal-tuning-curve modulations during the inter-trial-interval (ITI). As a result, the population faithfully represents the time that passes between trials. Because the length of the ITI contributes to learning-rate and the strength of the memory, as shown here for aversive-conditioning, our results suggest that it is mediated by amygdala-prefrontal networks themselves. This is the first demonstration of representation of long-time-scales (dozens of seconds) in primate amygdala networks, an order of magnitude longer than the typical duration used for the inter-stimulus-intervals (ISI). Therefore, this network can support formation and maintenance of temporal contingencies not only when they are cued explicitly by external stimuli, but also when embedded in the temporal structure of the task. Below we discuss the implications of this finding to valence-based learning.

Our finding of neural representation of time in BLA-dACC network could support several mechanisms that contribute to the effect of ITI duration on acquisition (learning) and conditioned response (CR, performance), as demonstrated across species and learning tasks^37–40^. Accordingly, we show here that the learning-rate and size of the CR were positively correlated with ITI duration^4–8^. The underlying mechanisms for this effect can consist of multiple parallel processes.

One appealing explanation of ITI related enhancement stems from the ratio between the ITI and the ISI (CS-US duration). Larger ratio is correlated with accelerated learning^8, 41^, in line with the larger informative power of the CS on the occurrence of the US^42^. Here we demonstrate one major component, that of time within the ITI. This is a more challenging aspect for a neural network and hence the main novelty in our study, because it requires representation of long time scales beyond that of persistent activity in single-cells (as widely demonstrated in CS-US trace-conditioning, CS-delayed-response memory tasks, and interval-reproduction tasks). Nevertheless, full examination of this specific hypothesis would require longer ISI to allow the CS response to decay, and it would be intriguing to explore neural implementation of ratio-dependent acquisition rate. Here, the ISI was short and predictable, and hence our finding of unbiased decoding-error along the ITI can enable a representation of duration and ratio.

Despite the elegance of this temporal account of ITI-duration and/or ITI-to-ISI ratio, several studies suggest that it might be insufficient. This is based on the observation that the proximity of an ITI-interfering event to the preceding^43^ or consecutive^44^ trial increased the effect on learning performance. As alternative, the opponent process theory postulates that after an aversive US, an opponent relief-like process is initiated that interferes with the learning and longer ITI allows for this process to terminate and for learning to be enhanced^37, 45^. Our finding that a larger proportion of BLA neurons are tuned to the middle of the ITI can support this, as the relief process evolves after the termination of the US yet decays towards the following trial, in line with safety signals reported in BLA neurons^46, 47^. Further, the opponent process can also be activated by CS-presentation alone as learning progresses, and reduced following extinction^45^. Accordingly, we found higher response magnitude during un-reinforced trials that diminish with extinction.

Another option is the formation of 2^nd^ order associations mediated by the amygdala^23–25, 48^, first between the CS and the US, and then between the ITI itself and the CS, or between the US-offset and the next CS (via the ITI ‘filler’^26^). However, we did not observe any behavioral changes during the ITI in expectation of the next trial, and we therefore think it is a less probable interpretation. Even if such 2^nd^ order association indeed takes place, it would require temporal activity as we report here that tiles and bridges the duration of the interval. Albeit, the ITI here was of varying random duration and hence the next CS cannot be faithfully predicted from US offset, yet the coming CS is a complete predictor for the US. Together with the lack of preparatory (CR-like) behavior during the ITI, this supports the interpretation that it is not a mediated association, but rather a representation of trial structure and/or statistics of the environment that is represented in amygdala networks^27^.

In addition, behavioral^43^ and neural^49–52^ studies suggest that rehearsal of memory can occur during the ITI, where additional processing contributes and consolidates learning^53^. Although it is not clear at this point how representation of time and rehearsal are related and integrated^54^, and evidence for neural rehearsal processes in the amygdala is still scarce, further characterization of BLA and dACC contribution to ITI related processing could shed light on these mechanisms and their putative complementary role.

The representation of long time scales on the order of dozens of seconds, and without an external cue as an anchor, is a new finding in the primate amygdala. One can consider several brain regions as candidates to supply this information to the amygdala. The striatum is one, shown to underlie and pace short time scales at hundreds of miliseconds^55^, but also to signal longer durations^56, 57^, yet mostly in specific dedicated time tasks^39, 58^. However, the striatum does not project directly to the BLA, yet the BLA does project to parts of the striatum^59–61^. In contrast, temporal information can reach the ACC via cortico and cortico-striatal loops^62–64^, and from there to the Amygdala^29^. Paralleling this, we observed a higher proportion of dACC neurons that scale their peak ITI modulation with the length of the recent ITI, on a trial-by-trial basis. Similarly, the representation of time can reach the amygdala via time-cells and event-integration in the hippocampus^65–67^. Whereas the exact functional role of such loops remains to be examined, it supports the idea that time perception is mediated by multiple overlapping neural systems which are flexibly engaged depending on the task requirements ^68^. In the current case, they can play a role in valence/emotional-based learning and support the demonstrated contribution of the ITI to memory strength.

An additional interpretation for the differential peaks of modulation observed in the two regions can come from functional considerations. The Amygdala was shown to signal safety^46, 47^, in accordance with its view as an absolute valence decoder ^69–71^. ACC neurons signal anxiety, threat, and risk^72, 73^, and importantly, the ACC plays a major role in error, context and attention^31–33, 74^. These functions are higher either right after or right before a trial. Therefore, the subtle distribution of roles might underlie the differential preference of ACC neurons to mediate information early and late in the interval, whereas amygdala neurons signal its middle.

Here, we focused on the reciprocal functional loop between the BLA and the dACC and found that together they span the complete ITI duration. Moreover, we observed potential transfer of temporal information in confined periods, early and late, by zero-lag synchronization between simultaneously recorded pairs of neurons. We identified three classes of pairwise interactions that synchronize early in the ITI, late, and more diffused population that spans the middle of it. This is in line with other types of information that transfer between the amygdala and the dACC at the single-cell level^19, 20, 36, 75–78^ and orchestrated by field-oscillations^36, 79, 80^.

These two seemingly different interpretations for the difference in peak distributions are not mutually exclusive and can serve both purposes. Namely, utilizing the tendency of one region to report a specific state aids to maintain the temporal marker for that state. This enable the network to use the underlying state of the animal to maintain additional information about the temporal structure across the network. Even if the immediate goal of the network is to represent the overall length of the ITI, it cannot do so by single-cell sustained activity, because of the high energetic demands for maintaining firing-rates. The tuning-curve-like approach allows this, and especially when distributed across regions.

To conclude, we find here that the activity of amygdala and dACC neurons hold information about long time-scales, and together support learning-rate and memory strength during aversive-learning. The results provide a new role for this network in maintaining time-scales to build the temporal and/or statistical structure of the environment, and provide a mechanism that explains how longer intervals promote learning and memory. In turn, it can also explain why under circumstances of multiple subsequent experiences, aversive memories are formed faster and exhibit strong-to-abnormal responses. Moreover, in addition to existing learning models that implicate this specific network in anxiety and fear-disorders^11, 35, 81, 82^, our findings suggest a new model for how deviations in representation and computation in this circuitry can lead to maladaptive and exaggerated behaviors.

## Acknowledgments

We thank Drs. Rita Perets and Yoav Kfir for scientific advice and help in experiments. We thank Drs. Eilat Kahana and Nir Samuel for help with medical and surgical procedures; Dr. Edna Furman-Haran, Nachum Stern and Fanny Attar for MRI procedures. This work was supported by ISF #26613 and ERC-2016-CoG #724910 grants to R. Paz.

## Author Contributions

R.P., U.L., A.H.T. designed the study; U.L. and Y.S. performed the experiments; R.P., U.L. A.H.T. analyzed the data; T.S. contributed to data analyses and interpretations. R.P., U.L., T.S., A.H.T. wrote the manuscript.

## Methods

### Animals

Two male macaca fascicularis (5–7 kg) were implanted with a recording chamber (27×27 mm) above the amygdala and cingulate cortex under deep anesthesia and aseptic conditions. All surgical and experimental procedures were approved and conducted in accordance with the regulations of the Weizmann Institute Animal Care and Use Committee (IACUC), following NIH regulations and with AAALAC accreditation. Food, water, and enrichments (e.g. fruits and play instruments) were available ad-libitum during the whole period, except before medical procedures that require deep anesthesia.

### Recordings

The monkeys were seated in a dark room and each day 3-6 microelectrodes (0.6-1.2 MΩ glass/narylene coated tungsten, Alpha Omega, Israel or We-sense, Israel) were lowered inside a metal guide (Gauge 25xxtw, OD:0.51mm, ID:0.41mm, Cadence inc. USA) into the brain using a head-tower and electrode-positioning-system (Alpha-Omega, Israel). The guide was lowered to penetrate the dura and stopped at 2-6mm in the cortex. The electrodes were then moved independently further into the amygdala or the dACC. Electrode signals were pre-amplified, 0.3Hz-6KHz band-pass filtered and sampled at 25Khz; and on-line spike sorting was performed using a template based algorithm (Alpha Lab Pro, Alpha Omega, Israel). Anatomical MRI scans were acquired before, during, and after the recording period and these scans were used to guide the positioning of the chamber on the skull at the surgery and to calibrate the positioning of the electrodes in the amygdala and the dACC.

### Behavior paradigm and stimuli

Detailed descriptions of the behavioral paradigm and our odor delivery system (olfactometer) have been previously reported^19^. Briefly, tone (900-2400Hz) presentation was triggered by respiration onsets and was followed, during the acquisition, with an odor that was released at the onset of the following respiration cycle, but not before 1 second from the tone presentation. Odors (propionic acid or banana-melon organic extracts diluted in mineral oil) were actively evacuated from the nasal mask by a vacuum hose and respirations were constantly monitored with two parallel connected pressure sensors (1/4” and 1” H2O pressure range, AllSensors).

During the habituation phase the tones (novel tones on each day) were presented 10 times, without any odor to follow. During the acquisition stage, the tones were presented 30 times each, and paired with aversive odors in all occasions. In some sessions, aversive and pleasant trials were intermingled in a pseudorandom order and paired with different tones (discrimination learning). In addition, in half of the sessions, the acquisition stage also included 10 presentations of unpaired (‘catch’) trials, i.e. tones that were not followed by an odor.

### Behavioral analysis

Learning dependent changes in the response to the CS were computed as area under the curve during 350 ms. following CS-onset compared with mean inhale volume in response to the same tone during habituation, as in previous work^19^.

Behavioral ITI data was aligned to the end of the ITI, namely to the beginning of the next trial (CS occurrence), because our goal was to examine if any anticipatory/preparatory behavior develops towards the next trial. To evaluate changes in inhale power during the ITI across trials we averaged the area under the curve of breathing data during 30 seconds of the ITI with a running window of 7 seconds and overlap of 2 seconds. To evaluate changes in inhale frequency (for 0.2 – 2 Hz., steps of 0.01 Hz.) we quantified spectrograms by obtaining the multitaper power spectral density estimation (Thomson multitaper method) for breathing data during 30 seconds of the ITI (7 sec windows with and 2 sec overlap).

### Neural data analysis

Neural activity during the ITI was aligned to the end of each trial, and taken until the end of the mean-ITI in each specific session, with exclusion of bins from trials with shorter ITI (to avoid sampling bias and CS-response contaminating the dataset). This is because our hypothesis was that neural activity signals the time during the ITI that passes from the previous trial (as the reliable way to estimate the duration of the ITI).

#### ITI-FR modulation

Spikes discharge that occurred between the US offset and the mean-ITI were binned into 6, 12, 20 or 40 bins and counted to produce an estimation of the ITI firing rate (ITI-FR). To assess the strength of modulation, spikes were shuffled 100 times and the mean ITI-FR was compared to the mean and standard deviation of the shuffled ITI-FR distribution. Data was taken only from bins that occurred before the ITI-mean, to avoid sampling bias. Up and down modulations were determined when the original spike-count at a specific bin exceed ±2 standard deviation of the shuffled spike count at the corresponding bin.

#### Acquisition of ITI response

ITI-FR was computed separately for habituation, acquisition and extinction ITIs. The ITI-FR of the acquisition and extinction was further separated into 3 and 2 equally sized (10 trials) sub-stages respectively. Each ITI-FR was then normalized according to its own shuffled dataset (as above). This normalization assures that the ITI-FR at all stages will be on the same scale even in cases that the neurons increase or decrease their firing rate along the session (i.e. non-stationary). Next, the root mean square (RMS) of the ITI-FR was derived and compared between the different stages. ANOVA test was employed to examine whether the RMS scores changed between the habituation and the acquisition. When the ANOVA test indicated that the RMS was modulated during the session, we further performed post-hoc t-test comparisons between the early, middle and late acquisition phases and the habituation. Neurons whose ANOVA test was significant and with one or more post-hoc comparisons that indicate that their RMS is increased during the acquisition stages were classified as acquisition responsive neurons.

#### Temporal Tuning-curve and peak-modulation (preferred-time)

Spikes were binned into 40 bins during the ITI, and the bin with the highest firing-rate (averaged over trials) was marked as the peak. This bin was then gradually joined by surrounding bins if they had a higher-than-baseline firing rate, to form the temporal tuning-curve. A one-way ANOVA (at p < 0.05) determined if the firing rate in these bins was higher than an equivalent number of bins from the rest of the ITI.

#### ITI correlations

Pearson correlations between the center-of-mass of the ITI-FR and the length of the previous ITI (duration) were computed. Correlation at p<0.05 were considered significant, and binomial test was employed to determine whether the percentage of units with significant correlation is higher than chance level (5%). Similarly, Peasron correlations were also computed between the RMS of individual ITI-FR during the acquisition and the length of the ITI at the previous trial.

#### ITI cross-correlation

Cross-correlations were computed between all pairs of amygdala-dACC that were recorded simultaneously and which firing rate exceed a mean of 1Hz during the ITI (to allow sufficient reliable statistical power). Cross correlations were computed in a three seconds window that was advanced in one second steps from the offset of the US until five (5) seconds before the mean ITI. In all cases, amygdala spikes were used as the reference for the occurrence of dACC spikes, and spike that occur up to 750 ms before or after an amygdala’s spike were included in our analysis. Next, spike times were binned into 20ms bins and counted. To evaluate whether the observed cross-correlations were significant we repeated the same procedure but this time, the dACC spikes were shuffled in two different manners: 1. *Within-ITI* shuffling, in which we shuffled spikes in each individual ITI, destroying their temporal resolution, but maintaining the overall firing rate in each individual ITI; and 2. across-*ITI* shuffling, in which spikes were shuffled between the different ITIs, but Maintaining the same epoch within the ITI. This procedure changed the overall firing rate of individual ITIs, but maintains the temporal structure of the ITI-FR. Cross-correlations were deemed significant only if they exceeded both criteria, with a further imposed requirement that they maintain significance for at least 15% of the total ITI duration.

#### Decoding time from neural activity

we used cross-validation and an optimal linear estimator (OLE). In each iteration, we pseudorandomly chose 25 trials from the acquisition and derived the optimal weights given the normalized firing rate in each bin of the actual time during the ITI (40 bins). These weights were then used to estimate/decode the time in each of the remaining 5 trials separately by using the real firing-rates. We then averaged across neurons. The process was repeated 100 times, and also as a function of N neurons pseudorandomly selected from the different sessions (see Fig.8). A population vector approach gave highly identical results.

## Supplementary Figures 1,2,3

**Supp.Fig.1.**
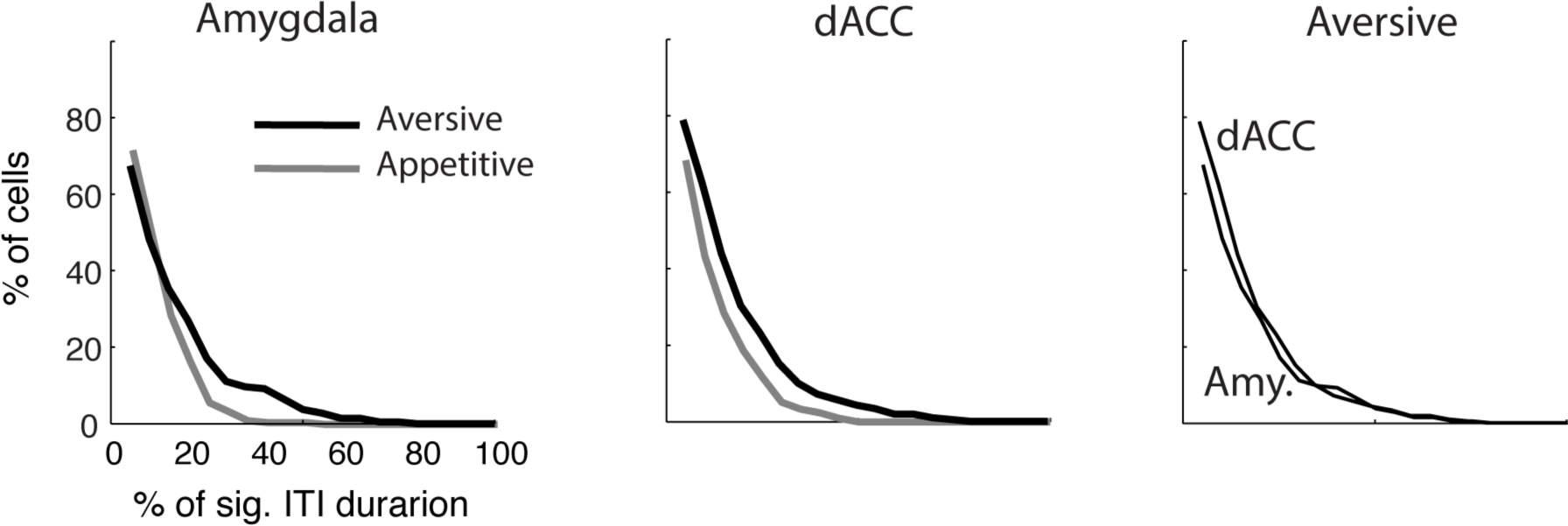
Proportions of cells with ITI-FR modulations. Similar to Fig.3E. Histograms of the proportion of cells (y-axis) that showed significant ITI-FR modulation (ITI-FR > 2 s.t.d of the shuffled FR) in a proportion of the ITI duration (x-axis). All histograms are significantly different than a null distribution obtained from shuffling (Fig.3E). Left: In the amygdala proportion of cells in aversive learning was slightly higher than in appetitive learning. Middle: Similarly in the dACC. Right: The proportion of cells in aversive learning was similar in the amygdala and the dACC.

**Supp.Fig.2.**
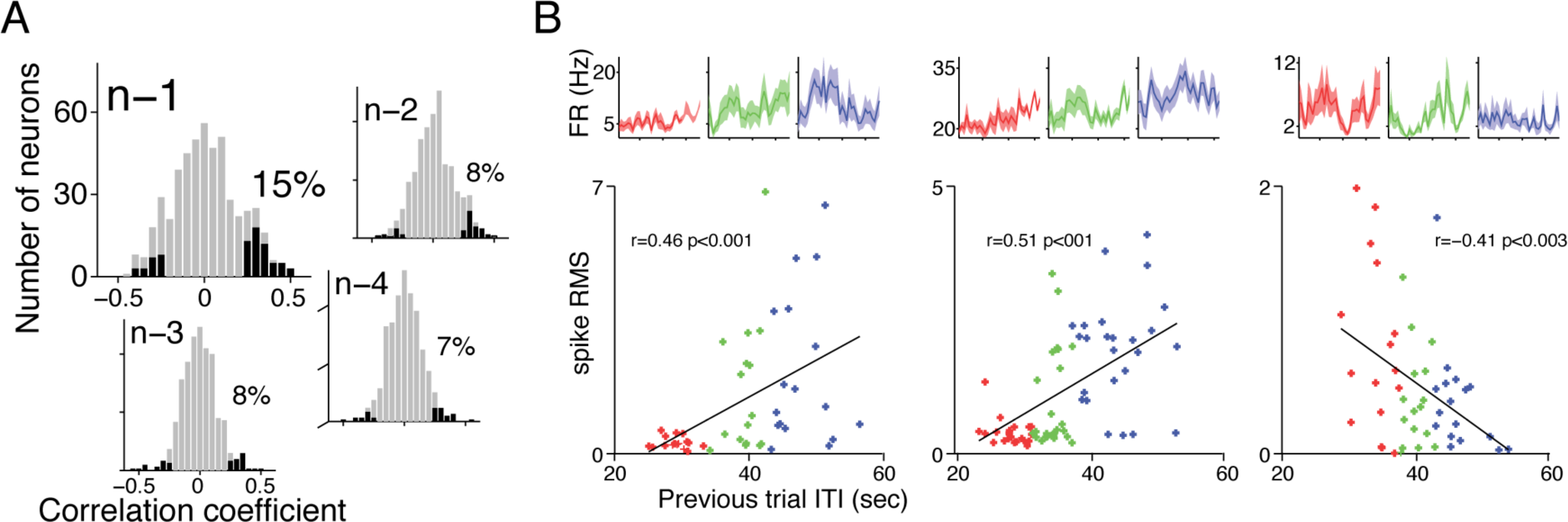
ITI duration affects the strength of peak-modulation of the next trial. Same presentation as in Fig.5. (A) The strength of the peak-modulation is correlated with the duration of the previous trial ITI in 15% of all cells (p<0.01, binomial test). Earlier ITIs (trials n-2/3/4) affect much less. (B) Three examples of cells in which the strength of modulation (quantified as excess in spikes as in Fig.4, y-axis) correlates with the ITI duration of the previous trial (x-axis). The PSTHs of the lower third (red), middle third (green) and upper third (blue) are shown on top.

**Supp.Fig.3.**
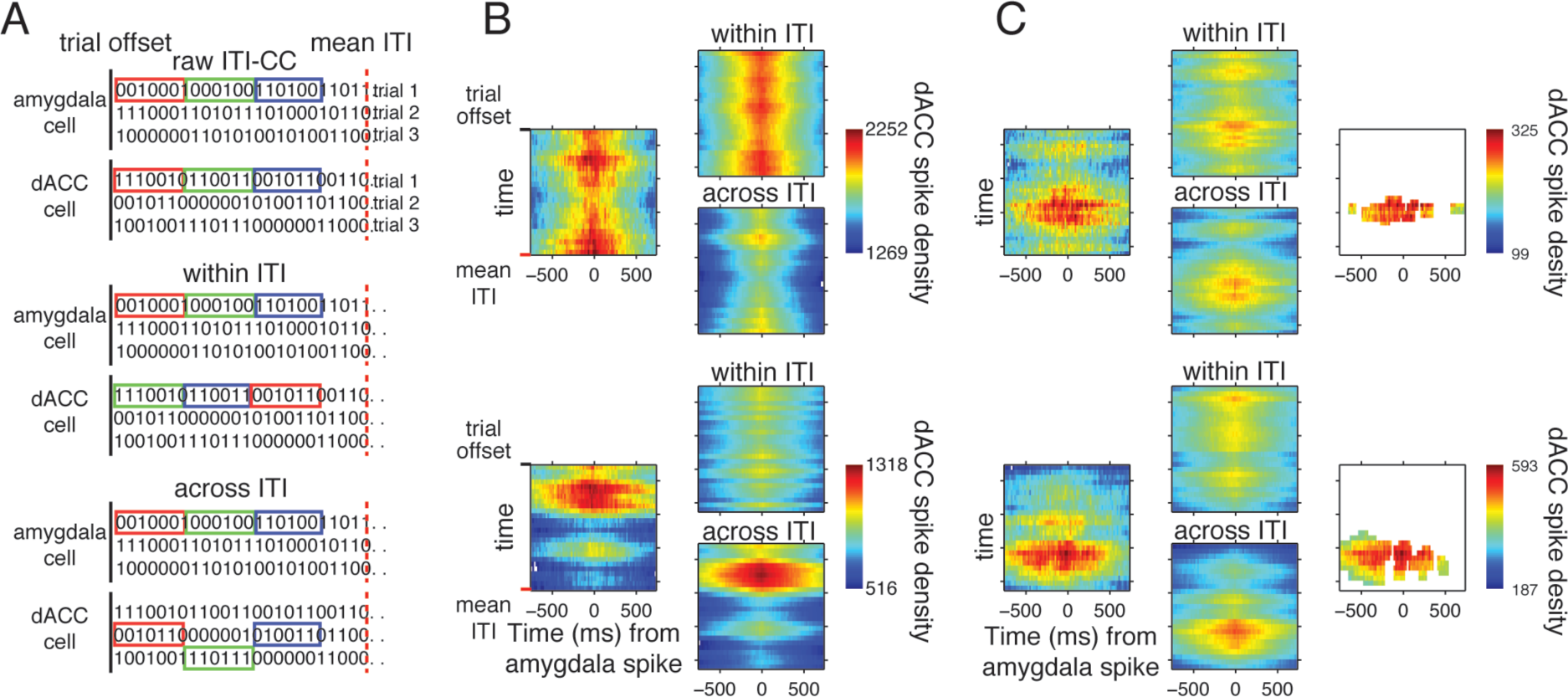
Quantifying amygdala-dACC pairwise cross-correlations (CC) along the ITI. Cross-correlations were computed between simultaneously recorded amygdala-dACC neurons in sliding 3-seconds segments of the ITI. All CCs are triggered on amygdala spike (at time zero). CCs were compared to shuffles to exclude changes resulting from covariations within and across ITIs. All maps show the time in the ITI on y-axis (top-to-bottom), and the CC itself as time from amygdala spike on the x-axis, colored by spike-density. (A) To identify the significant correlation bins we compared the raw CC (left in B,C) against two types of shuffled cross-correlations (A, and right maps in B,C). (1) within-ITI shuffle (A, middle column) exchange regions within each ITI, and removes correlations that stem from covariance in the FR along the ITI. (2) across-ITI shuffle (A, lower column) exchange the same temporal regions in the ITI across different trials, and removes correlations that stem from covariance in the FR along the trials. (B) Shown are two examples where only one shuffle, but not the other, account for the observed raw cross-correlations. Such CCs were not deemed significant. (C) Shown are two examples where neither of the shuffled cross correlations account for the raw cross correlation. Maps on the right show the bins deemed significant after comparison to the two shuffles.

